# Amazonian run-of-river dam reservoir impacts underestimated: Evidence from a Before-After Control-Impact study of freshwater turtle nesting areas

**DOI:** 10.1101/2021.07.29.454366

**Authors:** Andrea Bárcenas-García, Fernanda Michalski, James P. Gibbs, Darren Norris

## Abstract

1. Construction of hydropower dams is associated with negative impacts on biodiversity, yet there remains a lack of robust scientific evidence documenting the magnitude of these impacts particularly across highly biodiverse tropical waterways. Hydropower expansion is an increasing threat to the Endangered yellow-spotted river turtle (*Podocnemis unifilis*) across its tropical South American range.
2. Yellow-spotted river turtle nesting areas were monitored as an indicator of dry season river level changes following run-of-river dam reservoir filling. A Before-After-Control-Impact (BACI) study design was used with multi-year field campaigns monitoring turtle nesting areas upstream of the dam.
3. The cause and extent of changes in nesting areas were established using Generalized Additive Models. Nesting area density was evaluated in relation to: time (before versus after), treatment (control versus impact), time treatment interaction (BACI), distance to the dam and precipitation. The extent of changes was examined by comparing the proportion of nesting areas remaining during four years after reservoir filling.
4. Dam construction generated an immediate and apparently permanent dry season river level rise that extended more than 20 km beyond impact assessment limits. On average the density of nesting areas declined 69% (from 0.48 to 0.15 per km) across 33 km of river directly impacted by the dam. This loss was reflected in a significant BACI interaction. Nesting area density was not explained by seasonal precipitation.
5. Standardized monitoring of freshwater turtle nesting areas provided an effective means to quantify impacts of hydropower developments across biodiverse yet rapidly changing waterways. The negative impacts documented in this study should be preventable by mitigation actions including habitat creation and dry season flow regulation. Such measures would also likely benefit multiple species elsewhere in tropical rivers increasingly impacted by run-of-river dams.

## 1 INTRODUCTION

Mitigating dam impacts is a priority for the protection and restoration of freshwater biodiversity (Harper et al., 2021; Moran et al., 2018; Tickner et al., 2020). Hydropower developments are among the most significant threats to Amazonian freshwater ecosystems (Castello & Macedo, 2016; Latrubesse et al., 2017) and biodiversity (Dudgeon, 2019; Latrubesse et al., 2020; Vasconcelos et al., 2020). Although hydropower supplies the demand of basic human necessities, hydropower dams result in impacts that produce negative effects and feedbacks that modify freshwater ecosystem processes, functions and services (Dudgeon, 2019; Valiente-Banuet et al., 2015). These impacts generate changes not only to biological cycles and species ecological interactions (Latrubesse et al., 2020) but also create drastic alterations to local communities, livelihoods and cultures (Del Bene, Scheidel, & Temper, 2018; Fearnside, 2018). Whilst negative impacts have been widely suggested, more robust evidence is required to examine the sustainability of the increasing number of dams across Brazilian Amazonia (Rodrigues dos Santos, Michalski, & Norris, 2021; Tundisi et al., 2014).

The ecological structure and complexity of Amazonian freshwater ecosystems is a result of myriad factors operating across different temporal and spatial scales, including flood pulse (Junk & Wantzen, 2006; Melack & Coe, 2021) and the spatial-temporal heterogeneity of seasonal rainfall (Melack & Coe, 2021; Reis et al., 2019; Siddiqui et al., 2021). The structure, composition, functions and services of globally important Amazonian ecosystems are threatened due to expansion of hydropower developments. To date a total of 29 dams (operational capacity > 30 MW) have been installed across Brazilian Amazonia (SIGEL, 2021) and another 750 are planned or under construction (Vasconcelos et al., 2020). This increase is incentivized by increasing energy demand, national and regional politics and international investment (Castello, 2021; Fearnside, 2018; Gerlak et al., 2020). Although hydropower dams require Environmental Impact Assessments, those conducted in the Brazilian Amazonia are typically characterized by localized, unstandardized and short-term biodiversity monitoring (Fearnside, 2014, 2018; Pelicice & Castello, 2021).

There is no standardized typology of dam types or sizes, which can often generate confusion in the scientific literature. For example the International Commission of Large Dams (https://www.icold-cigb.org/, accessed 9 July 2021) defines a large dam as: “A dam with a height of 15 meters or greater from lowest foundation to crest or a dam between 5 meters and 15 meters impounding more than 3 million cubic meters”, but such definitions vary in different regional and national contexts. “Run-of-river” dams are widely considered to generate relatively less negative impacts compared with impoundment dams as the production of energy works with the natural river flow (Gaudard, Avanzi, & De Michele, 2018). Yet, “run-of-river” is not synonymous with small or low impact hydropower operations (Kuriqi et al., 2021), causing loss of longitudinal connectivity, habitat degradation, and simplification of the biota, which leads to increasing need for more economically and environmentally sustainable options such as in-stream turbines across Amazonia (Chaudhari et al., 2021).

Although adoption of newer technologies is likely to reduce environmental impacts in the future (Chaudhari et al., 2021; Moran et al., 2018), currently a lack of evidence limits the development of effective mitigating actions of Amazonian run-of-river hydropower dams (Gerlak et al., 2020; Gomes et al., 2020). Run-of-river dams may be constructed with or without reservoirs and range in size from small (< 1 MW) to large “mega-dams” such as the Belo Monte dam complex in Brazil (total installed capacity of 11,233 MW, dam heights 36 – 90 m). The use of natural hydrological flows reduces the energy production capacity of run-of-river dams during the dry season, which is a period when supplying the energy demands can influence dry season river flow and freshwater habitats (Gierszewski et al., 2020; Latrubesse et al., 2020). Indeed, although impacts of large hydropower developments could lead to species extinction (Turvey et al., 2010) and extirpation in impacted basins (Gomes et al., 2020; Santos et al., 2020) there remains a lack of information about the impacts and responses of freshwater vertebrates to hydropower dams (He et al., 2018).

Aquatic species, like freshwater turtles are directly impacted by dam construction (Le Duc et al., 2020; Stanford et al., 2020; Tucker, Guarino, & Priest, 2012). Although many studies focus on fishes there are few studies considering the diversity of Amazonian aquatic and semi-aquatic vertebrates (Rodrigues dos Santos, Michalski, & Norris, 2021). Turtles represent some of the largest and most threatened freshwater vertebrates (Stanford et al., 2020; Tickner et al., 2020) and are vulnerable to habitat changes because their life history (Shine & Iverson, 1995) and dependence on environmental factors for embryonic development, which limits the adaptive response for population recruitment against anthropic impacts (Quintana et al., 2019). Little is known about the impacts of hydropower dams on freshwater turtles at temporal and spatial scales relevant to develop effective management actions. As freshwater turtles are typically long-lived there is a need for long-term studies to detect life-history trends (Iverson, 1991; Shine & Iverson, 1995), but there are relatively few studies available documenting tropical freshwater turtle survival (Rachmansah, Norris, & Gibbs, 2020) and even fewer in relation to dam impacts (Bárcenas-García et al., 2021).

The Endangered (Rhodin et al., 2018) yellow-spotted river turtle (*Podocnemis unifilis*, Figure 1) is the most widespread of six *Podocnemis* species found in South America, all of which nest on dry land (Turtle Taxonomy Working Group, 2017). Although widespread, *P. unifilis* is increasingly threatened (Rhodin et al., 2018) and without effective conservation actions severe (≥50%) and rapid (<50 year) population losses were projected across 60% of the species 5.3 M km^2^ pan-Amazonian range (Norris et al., 2019). Studies emphasize the need to focus on adult survival for the conservation of freshwater turtles (Páez et al., 2015; Rachmansah, Norris, & Gibbs, 2020), but early stages (e.g., egg and hatchling) are also important for the conservation and recovery of freshwater turtles (Rachmansah, Norris, & Gibbs, 2020) and *P. unifilis* populations (Norris et al., 2019). Nesting patterns are useful to understand responses of chelonian populations (Iverson, 1991) to environmental disturbance because nesting is key for female turtles to guarantee their survival and sustain populations (Refsnider & Janzen, 2010). Changes in dry season flood pulse and/or flow rates, could therefore result in negative impacts on *Podocnemis* populations due to changes in nest site selection and hatchling success (Eisemberg et al., 2016; Norris, Michalski, & Gibbs, 2020).

**Figure 1.**
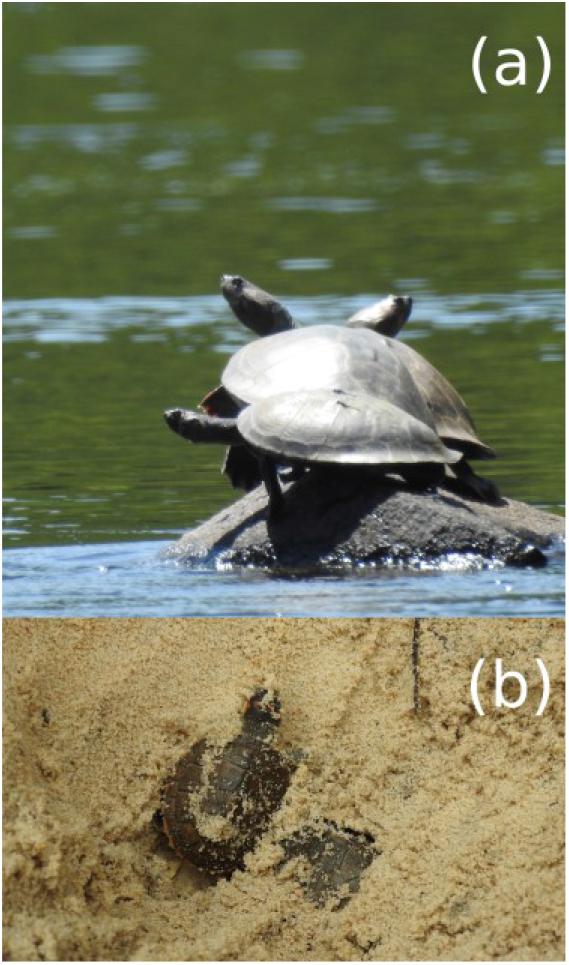
Study species. Adult and hatchlings of the yellow-spotted river turtle: (a) Adult females basking and (b) Hatchlings emerging from a nest laid in the sand of a riverside nesting area.

Freshwater turtle nesting areas are important for multiple Amazonian species not only for the turtles themselves (Campos-Silva et al., 2018). The seasonal reduction in river flow exposes river banks and islands, creating seasonally ephemeral habitats that represent essential resources for both freshwater and terrestrial biodiversity across Amazonia (Castello, 2021; Junk & Wantzen, 2006; Melack & Coe, 2021). These seasonal habitats are used by diverse oviparous wildlife (Campos-Silva et al., 2018; Campos-Silva et al., 2021) and their predators (da Costa Reis et al., 2021), and can also facilitate movements e.g. as stepping-stones to cross rivers (Mamalis et al., 2018), and provide locations for reproductive/territorial behaviours (Michalski et al., 2021).

In the present study *P. unifilis* nesting areas were monitored as a biologically relevant indicator of river level changes following run-of-river dam reservoir filling. It could be expected that after reservoir filling the shorelines in the area of dam influence may develop characteristics more favourable to nesting by *P. unifilis*. The objective was to expand on previous results from the first year after construction of a large run-of river dam (Norris, Michalski, & Gibbs, 2018a), comparing annual numbers of turtle nesting areas to examine if upstream patterns in the submersion of nesting areas remained consistent four years after reservoir filling and to test the influence of potentially confounding variables (seasonal precipitation). This comparison across multiple years provides for the first time robust causal evidence documenting the spatial and temporal extent of impacts of a run-of-river dam reservoir on an Endangered vertebrate species in the Amazon. The study documents immediate and apparently permanent dry season changes beyond established limits with implications for global efforts to monitor free flowing rivers (Grill et al., 2019). The findings are used to suggest important mitigation actions that are likely to benefit multiple species in highly diverse Amazonian rivers increasingly impacted by run-of-river dams.

## 2 MATERIALS & METHODS

### 2.1 Study area

This study was conducted in the Araguari river basin, eastern equatorial Amazonia, in the central region of the Brazilian State of Amapá (Figure 2). The basin includes rivers classified as “clear water” (Junk et al., 2015). The Araguari River has an extension of 560 km (Ziesler & Ardizzone, 1979) and rises in the Guiana Shield at the base of the Tumucumaque uplands, bifurcates and discharges into the Amazon River and the Atlantic Ocean. The region’s climate is humid tropical [“Am” Tropical monsoon (Kottek et al., 2006)], with a dry season from September to November (<150 mm of monthly rain) and the rainy season (>300 mm of monthly rain) from February to April (Supporting Information S1).

**Figure 2.**
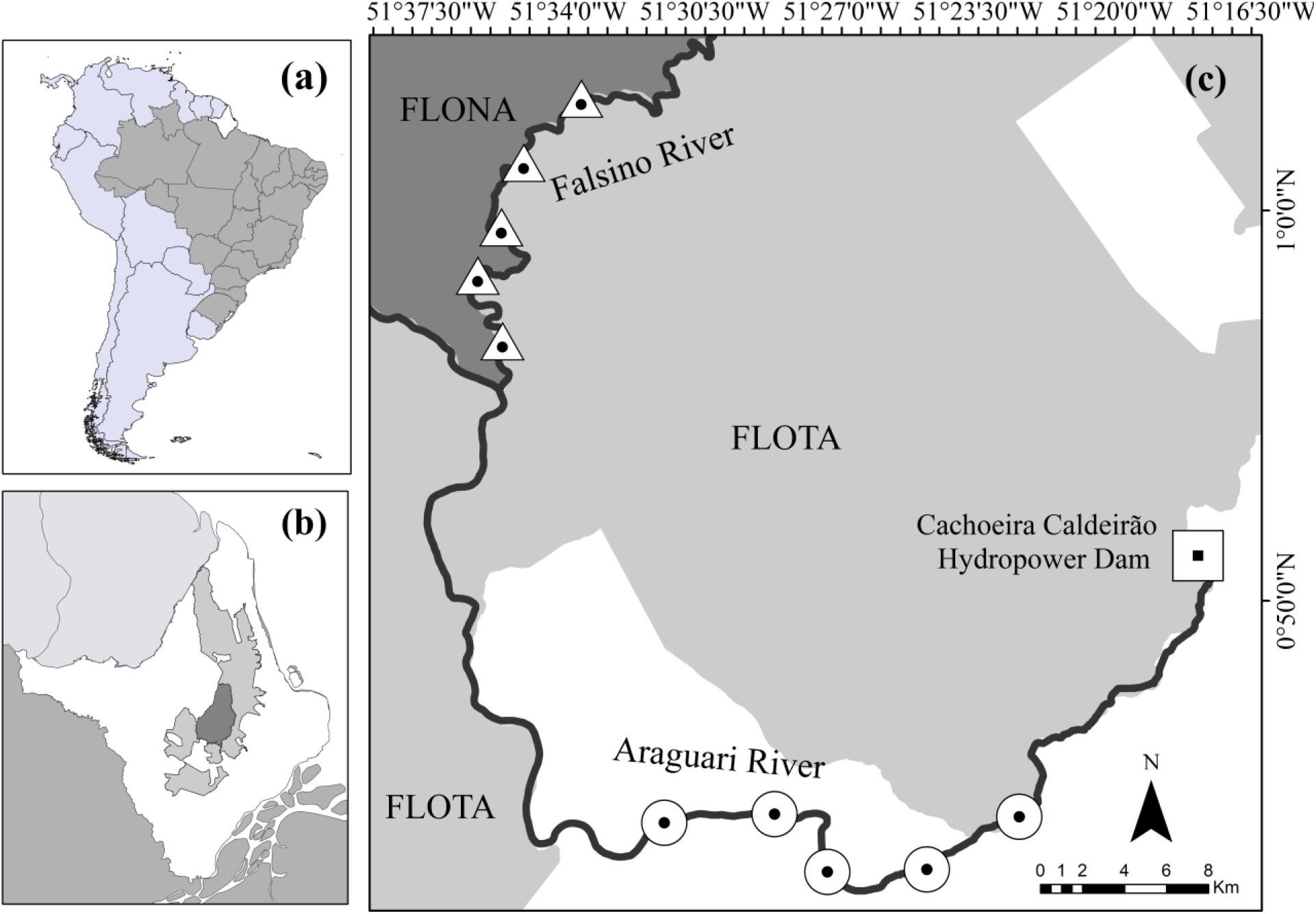
Study area. (a) State of Amapá in Brazil. (b) Location of the study river (black line) within Amapá. (c) Location of the Impact (circles) and Control (triangles) subzones upstream of the Cachoeira Caldeirão hydropower dam (square). Hashed shading is the area of direct impact identified in the Environmental Impact Assessment. Grey shaded polygons show the protected areas: Amapá National Forest (“Floresta Nacional do Amapá” - FLONA) and the Amapá State Forest (“Floresta Estadual do Amapá” - FLOTA).

There are currently three dams with a combined installed capacity of 549 MW (78, 252 and 219 MW) along an 18 km stretch of the Araguari River (INESC, 2021; SIGEL, 2021). Coaracy Nunes was the first dam installed along the Araguari River in 1975 and was also the first dam in the Brazilian Amazon. The present study focuses on the 219 MW Cachoeira Caldeirão dam (51°17’W, 0°51’N), which became operational in 2016 and is the most recent and most upstream dam in the Araguari river. Cachoeira Caldeirão is a large run-of-river dam, with a height of 20.6 m and length of 730 m (EDP-Brasil, 2021b). It was installed and is currently operated by the Empresa de Energia Cachoeira Caldeirão S.A, which is a joint venture between China Three Gorges Brasil and Energias de Portugal-Brasil S.A (EDP-Brasil, 2021a). Upstream of the dam, the Araguari River flows between two sustainable-use conservation areas (Figure 2): Amapá National Forest (“Floresta Nacional do Amapá” - FLONA) and the Amapá State Forest (“Floresta Estadual do Amapá” - FLOTA).

### 2.2 Nesting-area surveys

Monthly boat surveys were conducted to locate nesting areas (including beaches, sandbars, exposed river bank margins) from September to December, which corresponds to the nesting season of *P.unifilis* in the study area (Norris, Michalski, & Gibbs, 2018a). The majority of *P. unifilis* nesting studies come from larger rivers where nesting areas are large (> 1 ha) and likely to persist across decades (Campos-Silva et al., 2018; Escalona & Fa, 1998). In contrast the nesting areas upstream of the Cachoeira Caldeirão hydropower dam are typically smaller (0.03 ha on average (Norris, Michalski, & Gibbs, 2018a), Figure 3) and can be ephemeral between and within years due natural changes in flow and river levels (Quintana et al., 2019).

**Figure 3.**
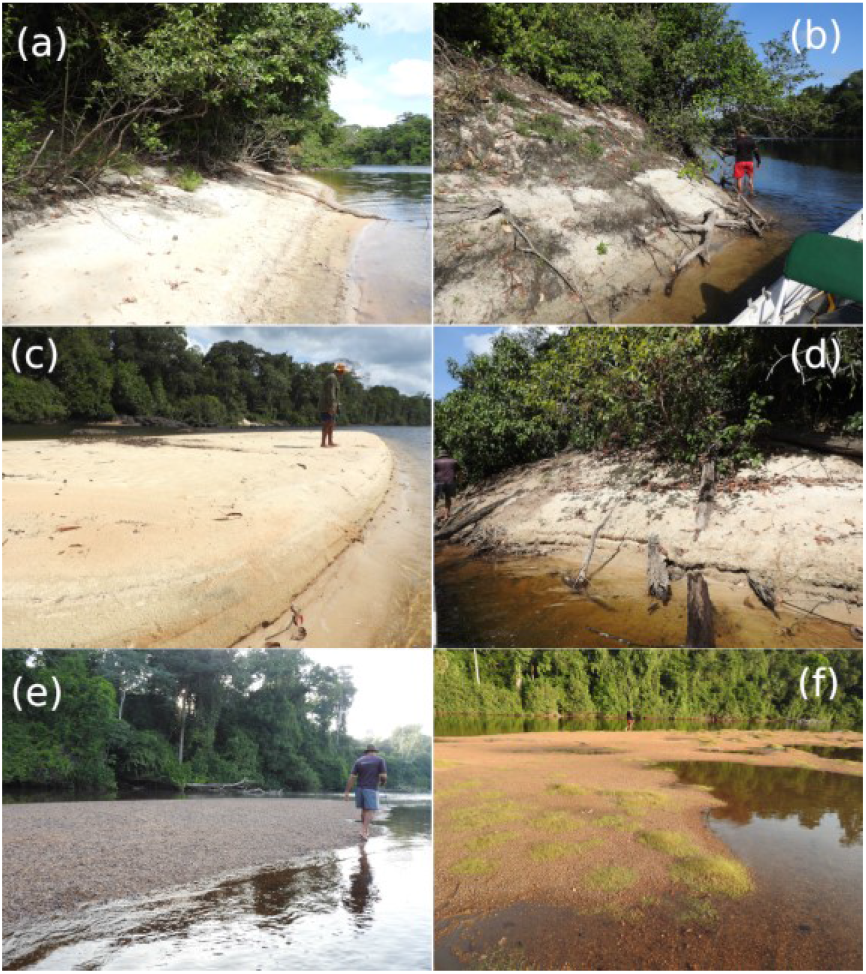
Nesting-areas. Representative yellow-spotted river turtle (*Podocnemis unifilis*) nesting areas examples showing (a, b): potential nesting-areas with suitable habitat conditions for nesting, but no nests (i.e., locations that females could use for nesting but nests were not found); (c, d): actual nesting-areas (i.e., locations where females actually nested), and (e, f): unsuitable habitat areas for nesting (i.e., where substrate conditions were not appropriate for females to nest). Reproduced from Norris, Michalski, and Gibbs (2018a).

Surveys were carried out upstream of the Cachoeira Caldeirão hydropower dam before (2011 and 2015) and after (2016, 2017, 2018, 2019) reservoir filling. Sampling and data collection in 2019 followed that from previous years (2011, 2015, 2016, 2017 and 2018) as described previously (Michalski et al., 2020; Norris, Michalski, & Gibbs, 2018a; Norris, Michalski, & Gibbs, 2020; Quintana et al., 2019). Here the standardised sampling methods are briefly summarised. Prior to dam construction in 2011 two separate 33 km river sections were surveyed for nesting areas, the first impact zone starting approximately 13 km upstream from the dam, the second control zone starting approximately 72 km upstream from the dam. From 2015 a continuous stretch of river was surveyed, thereby completing the gap between impact and control zones initially monitored in 2011 (Norris, Michalski, & Gibbs, 2018a).

During the monthly surveys areas suitable for turtle nesting (Figure 3) were located and identified based on the following features: areas >1m^2^, with exposed sand/gravel substrates, raised above river level, without being waterlogged to a depth of 15 cm (Norris, Michalski, & Gibbs, 2018a). For each nesting area it was recorded whether the surface was exposed or submerged below water. Nesting-areas that were flooded subsequent to exposure/nesting were identified as flooded and not included as “submerged” i.e. submerged includes only those nesting areas that were never exposed during the nesting season (September to December). Actual nesting areas were identified prior to reservoir filling as those where females nested in 2015 and/or in the previous five years [2010 – 2014, (Norris, Michalski, & Gibbs, 2018a)]; post-reservoir filling the actual areas included those where females nested in the current nesting season and/or during the previous five years.

To confirm the presence of turtle nests, searches were conducted by a team of three observers. One member was a local resident with over 30 years of knowledge of nesting areas. Searches were made at a standardized walking speed (mean 0.8 km per hour, range 0.2-1.3) with time spent searching areas ranged from 10 to 97 minutes depending on the size of the area (Norris, Michalski, & Gibbs, 2018a; Quintana et al., 2019). Nests were located by following turtle tracks on the substrates (sand/gravel) and systematic substrate searches with a wooden stick (Norris, Michalski, & Gibbs, 2018a; Quintana et al., 2019). Nests that had been depredated were identified by the presence of broken eggshells outside the nest, disturbed/uncovered nests and the presence of excavation marks (Escalona & Fa, 1998). The presence of all nests was considered including those depredated by humans or wildlife.

As there was potential for new nesting areas to emerge or to be lost due to natural changes in river flow both actual and potential nesting areas were monitored to understand the extent of dam impacts. Potential areas were those where environmental conditions matched those described in the literature and/or those found at the nesting areas from 2011, but without evidence or history of nest sites (Norris, Michalski, & Gibbs, 2018a). A nesting area could be either potential or actual during the nesting period. That is, if a potential area was identified and in a subsequent visit a nest was found then the area was counted only as actual during the nesting season.

### 2.3 Explanatory variables

Distance from dam and precipitation were used to explain temporal and spatial patterns in nesting area density. Distance to the hydropower dam was calculated along the river network using the riverdist package (Tyers, 2020) in R (R Core Team, 2020). Rainfall is an important determinant of annual and seasonal variation in Amazonian river flow rates (Melack & Coe, 2021; Siddiqui et al., 2021), as such elevated precipitation during the months preceding nesting could result in nesting areas remaining submerged (Eisemberg et al., 2016). The total and standard deviation of precipitation (PPT) during each of the three months (July, August and September) prior to the peak of the nesting season in the six monitored years was calculated (Supporting Information S1). Precipitation data was obtained from a weather station (Serra do Navio, id: 8052000, 52°00’W, 0°53’N), located approximately 115 km upstream from the Cachoeira Caldeirão dam; data available in the virtual database of the Brazilian National Water Agency (“Agência Nacional de Águas”: http://www.snirh.gov.br/hidroweb/serieshistoricas).

### 2.4 Data Analysis

Two different datasets were analysed, one to establish the cause of changes in *P. unifilis* nesting area density (Supporting Information S2) and another to quantify the extent of changes (Supporting Information S3).

#### 2.4.1 Cause of changes

To establish the cause of changes the response of actual nesting area density was modelled using Generalized Additive Models [GAMs, (Marra & Wood, 2011; Pedersen et al., 2019)]. To represent the Before-After-Control-Impact (BACI) sampling design (Smith, 2006; Underwood, 1993), the response of actual nesting areas per km was modelled by including two fixed factors, (i) before-after and (ii) control-impact and their interaction, with a significant interaction indicating differences between the impacted compared with the control zone associated with reservoir filling (Underwood, 1993). In this study, the impacted zone (Figure 2) corresponds to 33 km of the Araguari River, within the upstream area of direct dam impact [as defined by the Environmental Impact Assessment (Ecotumucumaque, 2013; EDP-Brasil, 2021b)]. A 33 km stretch of the Falsino River 72 - 105 km upstream of the dam (Figure 2), which corresponds to a region with little anthropogenic disturbance was established as the control zone (Norris, Michalski, & Gibbs, 2018a). For the GAM analysis each zone (control, impact) was divided into five subzones with an equal length of approximately 5.3 km (Supporting Information S2).

Spatial autocorrelation was quantified using Mantel test and semi-variogram (Dale & Fortin, 2014), which confirmed that each subzone could be considered as a spatially independent sample unit (Supporting Information 4). An information theoretic model selection approach (Burnham & Anderson, 2002) was applied to evaluate the alternative working hypothesis (Table 1). A set of seven models represented the working hypotheses to explain the patterns in nesting area density (Table 1). The five explanatory variables (total precipitation, standard deviation precipitation, distance to the hydropower dam, before-after, control-impact and the BACI interaction) were modelled separately to test their relationship with nesting areas per kilometre. All models included subzone and year as random effects to account for unexplained variation that was not part of the sampling design and enable estimation of the mean and variance of the global distribution of explanatory variables (Pedersen et al., 2019). An intercept only null model (without predictors) and a random only model (subzone and year only) were also included for comparisons. GAMs were run with Tweedie error distribution to examine which predictors described changes in nesting area density. Tweedie is a probability distribution family that includes the purely continuous normal and gamma distributions, the discrete Poisson distribution, and the class of compound Poisson gamma distributions (Jorgensen, 1997). All GAM smoothing parameters were estimated using Restricted Maximum Likelihood (REML).

**Table 1.** Hypotheses and variables used to explain the impacts of a run-of-river dam on yellow-spotted river turtle (*Podocnemis unifilis*) nesting areas in the eastern Brazilian Amazon.

| Category | Variable | Hypothesis | Description |
| --- | --- | --- | --- |
| Null | Intercept only |  | Response variable modelled without predictors. |
| Null | Random only |  | Includes years and subzone as random effects. |
| Environmental | Precipitation | Precipitation contributes to increase the river level, modifying the availability of nesting areas for *P. unifilis*. | Continuous. Total and variation (SD) in monthly precipitation (July to September) before peak *P. unifilis* nesting. |
| Anthropic | Distance to the hydropower dam | Areas closer to the dam have a higher negative impact. | Continuous. Distance in kilometres from the hydropower dam. |
| Anthropic | Treatment (Control versus Impact) | The impacted zone will have reduced nesting areas compared with the control zone. | Categorical with two levels. Impact Zone (Araguari River) and Control Zone (Falsino River). |
| Anthropic | Time (Before-After) | After four years operating, the dam has affected the nesting patterns for the *P. unifilis*. | Categorical with two levels. Years before (2011 and 2015) and after (2016, 2017, 2018 and 2019) reservoir filling. |
| Anthropic | BACI (Before-After-Control-Impact) | Significant interaction effect means that differences in nesting patterns are a direct result of dam impact. | Categorical interaction. The river section and period variables interact to represent the Before-After-Control-Impact (BACI) sampling design. |

To identify support for the seven alternative candidate models two Information Criteria were compared (Aho, Derryberry, & Peterson, 2014; Burnham & Anderson, 2004): Akaike Information Criterion corrected for small-sample sizes (AICc) and Bayesian Information Criterion (BIC). Model weights were calculated representing the likelihood of a given model, with evidence ratios constructed as the ratio of weights for the alternative models being compared (Burnham & Anderson, 2002). Model selection was followed by variable selection to establish the strength and direction of individual variables. Variable selection was based on a full model (including distance, precipitation and random variables) that was evaluated using a double penalty approach to determine which variables had the strongest effect on the response variable (Marra & Wood, 2011). As preliminary analyses suggested differences following reservoir filling, an interaction with the Before-After factor was included in each of the smooth terms for distance to dam, total precipitation and SD precipitation in this variable selection analysis.

#### 2.4.2 Extent of changes

From 2015 a continuous stretch of river was surveyed therefore the extent of changes in nesting areas was examined by comparing the number of nesting areas remaining in 6.6 km subzones (n = 15) during the four years after reservoir filling (2016, 2017, 2018 and 2019). From 2016 to 2020 the proportion of remaining nesting areas in each of the 15 subzones was calculated. The proportion of remaining nesting areas from 2016 to 2020 was then related to the distance of the subzone midpoint from the dam using a GAM with quasibinomial error family. All the analyses were performed in R 4.0.1(R Core Team, 2020), with the “mgcv” package to generate GAMs (Wood, 2020) and the “MuMIn” package (Barton, 2020) used for model selection.

## 3 RESULTS

Nesting area density decreased following dam construction, with the average density dropping from 0.48 areas per km before to 0.15 areas per km after reservoir filling at the Impact zone (Figure 4). Although Control subzones tended to have more nesting areas (Figure 4), there was variation in values between subzones and years, such that prior to reservoir filling there was no significant difference in mean nesting area density between Control and Impact zones (GAM, *P* = 0.279). In the first year following reservoir filling nesting area density declined to 0.06 nesting areas per kilometre in the impacted zone, with mean values remaining low in all following years (Figure 4). Although there was still some variation between years and subzones (values in the Impact subzones ranged from 0 to 0.91 nesting areas per km) there was a significant difference (GAM, *P* = 0.0061) in the average of nesting area density between Impact and Control zones after reservoir filling. Indeed, following reservoir filling the observed Control zone nesting area density was five times greater than that found in the Impact zone (0.74 and 0.15 areas per km, Control and Impact zones respectively).

**Figure 4.**
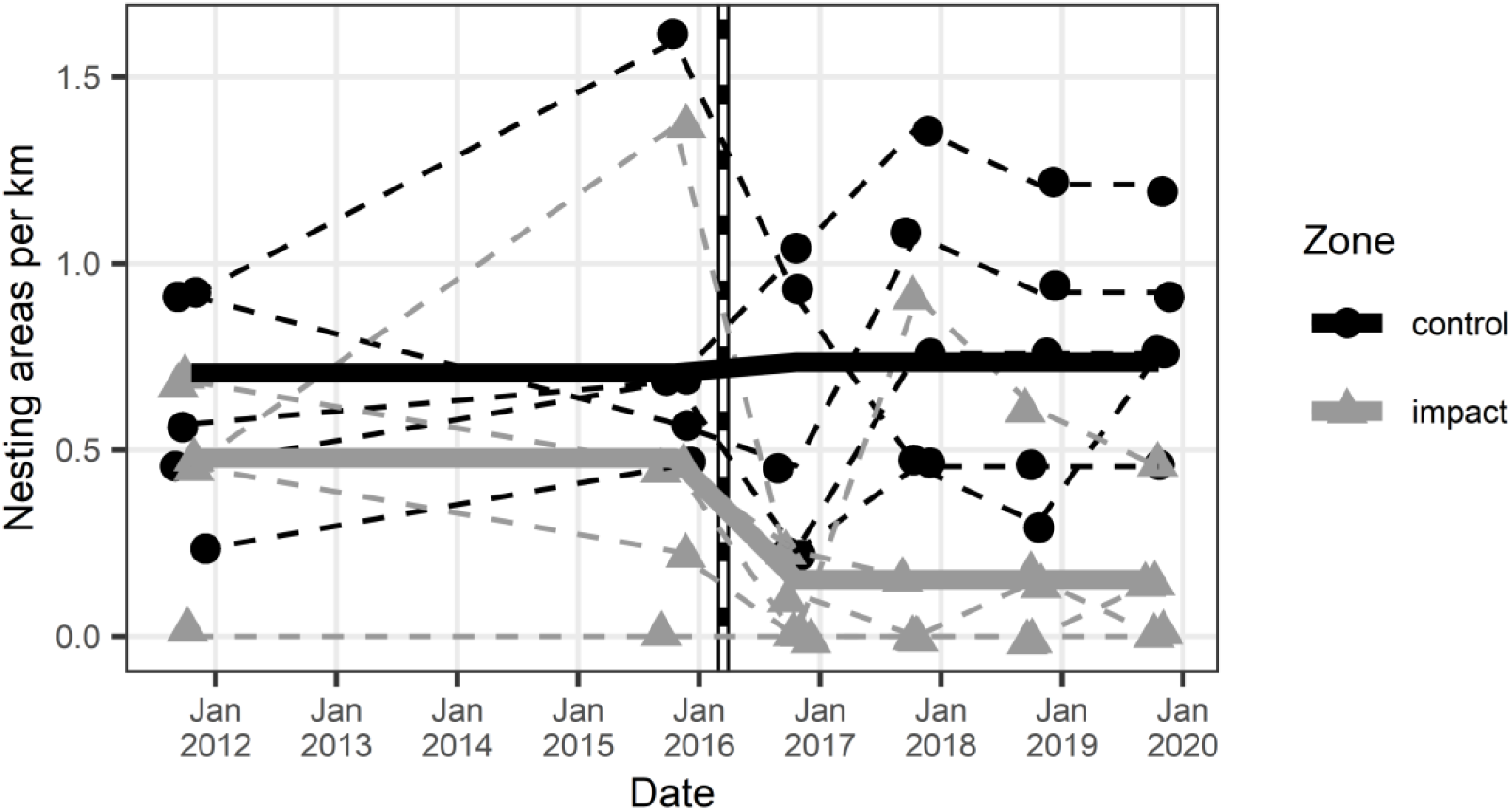
Nesting-areas before and after run-of-river dam construction. Observed patterns in yellow-spotted river turtle nesting areas in the eastern Brazilian Amazon. Density of *P. unifilis* nesting areas in nesting seasons before (2011, 2015) and after (2016 to 2019) dam reservoir filling in two zones (control and impact). Points show the number of nesting areas per kilometre in five 6.6 km river subzones in each of the impact (triangles) and control (circles) zones. Dashed lines join the yearly observations for each 6.6 km river subzone. Solid lines (black and grey) are predictions from Generalized Additive Model. Vertical dashed line represents when the dam reservoir was filled.

The model selection analysis indicated that the best model explaining patterns in nesting area density was the BACI (Table 2). All other models were only weakly supported (Table 2, ΔAICc > 4 and *wi* = 0.0 for models ranked by both AICc and BIC). The model including only random effects performed better compared with models including precipitation or distance to dam, with the random model having greater deviance explained and lower information criteria values (Table 2).

**Table 2.** Explaining patterns in yellow-spotted river turtle nesting in the eastern Brazilian Amazon. Generalized Additive Models were used to explain patterns in turtle nesting areas per km of river from 2011 – 2019. Models ranked according to an Information criteria approach using AICc (Akaike Information Criterion corrected for small sample sizes) and BIC (Bayesian Informative Criterion). K= number of parameters, Δ =delta for respective criterion values, and *wi* = the model selection weights. Models ordered by AICc values. Model construction including the following variables: time (“BA” before versus after dam construction), treatment (“CI” control versus impact zones), time treatment interaction (“BA:CI”), distance to dam (“Dist. to dam”), total rainfall (“PPTtotal”), standard deviation of rainfall (“PPTsd”), survey year (“Year”), sample location (“Subzone”).

| Hypothesis tested | Model construction | K | $R^2_{adj}$ | Dev. Exp. (%) | AICc | | | BIC | | |
| --- | --- | --- | --- | --- | --- | --- | --- | --- | --- | --- |
| | | | | | AICc | $\Delta$ | $w_i$ | BIC | $\Delta$ | $w_i$ |
| BACI | $Y \sim BA:CI + s(Year^\dagger) + s(Subzone^\dagger)$ | 12 | 0.5 | 55.0 | 64.7 | 0.0 | 0.8 | 83.6 | 0.0 | 0.7 |
| Before-After | $Y \sim BA + s(Year^\dagger) + s(Subzone^\dagger)$ | 12 | 0.4 | 50.1 | 69.2 | 4.5 | 0.0 | 88.0 | 4.4 | 0.0 |
| Random | $Y \sim s(Year^\dagger) + s(Subzone^\dagger)$ | 12 | 0.4 | 48.0 | 71.0 | 6.2 | 0.0 | 89.7 | 6.0 | 0.0 |
| Control-Impact | $Y \sim CI + s(Year^\dagger) + s(Subzone^\dagger)$ | 11 | 0.4 | 44.6 | 71.3 | 6.5 | 0.0 | 89.0 | 5.3 | 0.0 |
| Distance to dam | $Y \sim \text{Dist. to dam} + s(Year^\dagger) + s(Subzone^\dagger)$ | 11 | 0.4 | 46.2 | 71.5 | 6.8 | 0.0 | 89.8 | 6.2 | 0.0 |
| Precipitation | $Y \sim s(PPTtotal) + s(PPTsd) + s(Year^\dagger) + s(Subzone^\dagger)$ | 12 | 0.4 | 49.0 | 71.6 | 6.8 | 0.0 | 90.7 | 7.0 | 0.0 |
| Intercept | $Y \sim 1$ | 3 | 0.0 | 0.0 | 95.3 | 30.6 | 0.0 | 101.2 | 17.5 | 0.0 |

$s$ : Non-parametric smooth terms
† Random effects
$R^2_{adj}$ : Adjusted R squared for the model
Dev. Exp. (%): Percent of total deviance explained

A closer examination of the variables using a variable selection analysis indicated that distance to the dam was only significant after reservoir filling (Table 3, *P* = 0.0022, Supporting Information 5). There was no relationship between nesting area density and total precipitation. Prior to reservoir filling SD precipitation was weakly related to nesting area density (Table 3, P = 0.0147). Following reservoir filling there was no relationship between nesting area density and either of the measures of precipitation (Table 3, P = 0.1769 and 0.8809, total and SD respectively).

**Table 3.** Variable selection analysis. Smooth terms from a Generalized Additive Model used to explain patterns in yellow-spotted river turtle nesting areas per km of river from 2011 – 2019. Model construction using the following variables distance to dam, total rainfall (“PPTtotal”), standard deviation of rainfall (“PPTsd”), survey year (“Year”), sample location (“Subzone”).

| Non-parametric<br>smooth terms | EDF |  | P-value |  |
| --- | --- | --- | --- | --- |
|  | Before | After | Before | After |
| Distance to the dam | 0.00 | 0.90 | 0.5951 | 0.0022 |
| PPTtotal | 0.00 | 0.48 | 0.7239 | 0.1769 |
| PPTsd | 0.84 | 0.00 | 0.0147 | 0.8809 |
| Year<sup>†</sup> | 0.00 |  | 0.7801 |  |
| Subzone<sup>†</sup> | 6.47 |  | 0.0049 |  |
| R<sup>2</sup><sub>adj</sub> | 0.4 |  |  |  |
| Dev. Exp. (%) | 55.8 |  |  |  |

EDF: Estimated degrees of freedom for the model terms. Values close to zero indicate no relationship with the response, close to 1 may suggest a linear relationship and values greater than 1 suggest a non-linear relationship.
<sup>†</sup> Random effects
R<sup>2</sup><sub>adj</sub>: Adjusted R squared from the model
Dev. Exp. (%): Percent of total deviance explained

Comparison of nesting areas post reservoir filling showed significant non-linear increase in the proportion of remaining areas further from the dam in all four years post-reservoir filling (GAM, EDF = 8.61, P < 0.0001, Figure 5). Impacts were at least 20 km beyond those established by the Environmental Impact Assessment and 30 km beyond those used to establish free flowing river status (Figure 5). Within 50 km of the dam [official reservoir extension is 52 km (EDP-Brasil, 2021b)] between 76 and 100% of nesting areas were submerged following reservoir filling, but more areas remained further from the dam (20 to 76% from 49 to 72 km). There was much less variation in the control zone, with the proportion of remaining areas ranging from 87 to 100% in the five subzones beyond 75 km from the dam (Figure 5). Four years after reservoir filling an average of 25% of nesting areas were submersed 75 km upstream of the dam some 20 km beyond recognized limits. This loss was at least double that experienced in the five control subzones (mean 1 – 9% of nesting areas submersed, Figure 5).

**Figure 5.**
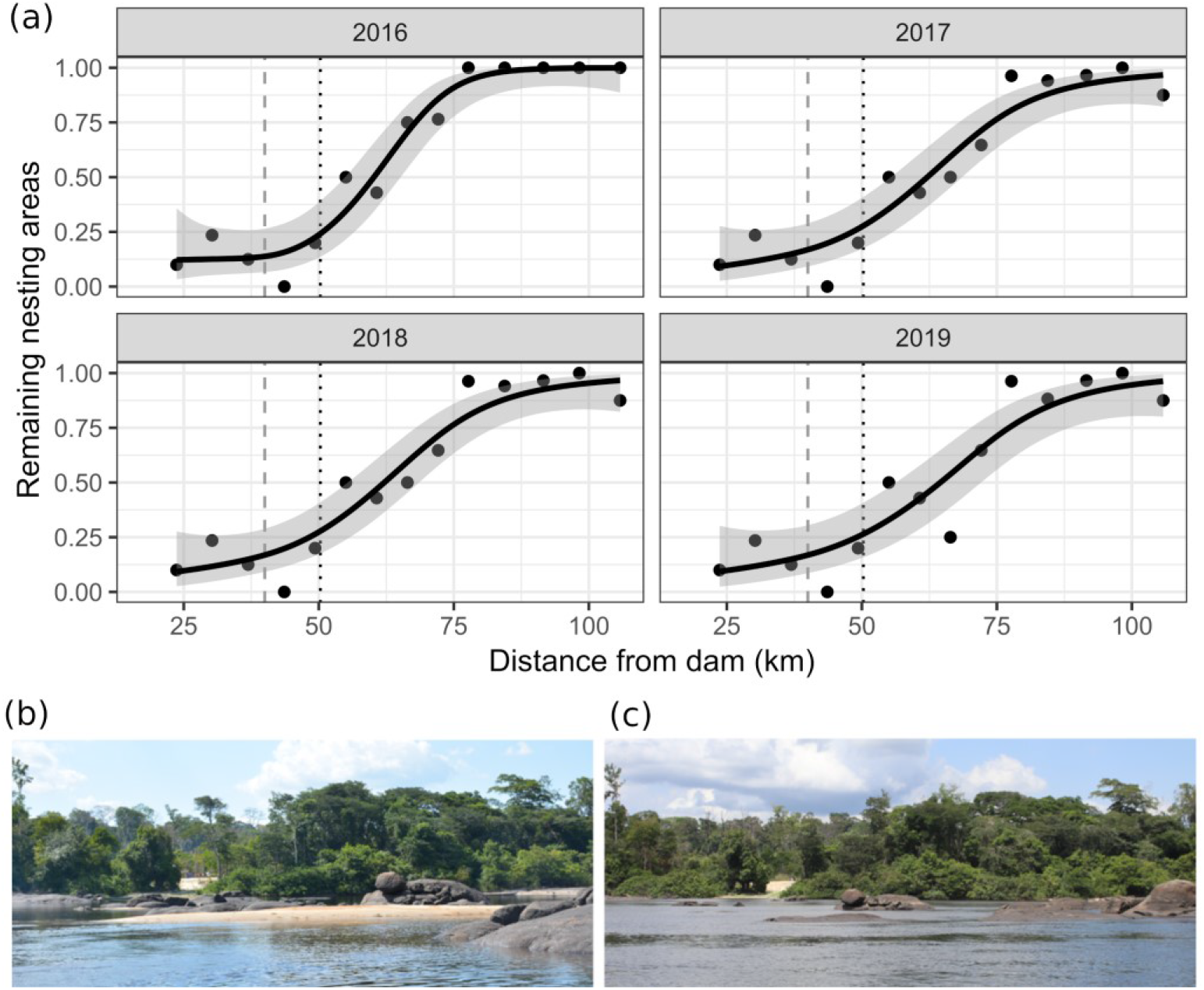
Extent of reservoir impacts. Remaining nesting areas following reservoir formation at the run-of-river Cachoeira Caldeirão Dam, Amapá, Brazil. (a) The proportion of nesting areas remaining in four years following reservoir formation was calculated along 15 equally spaced river sections (graphs generated excluding one section without nesting areas). Dashed vertical line represents the limit (upstream river confluence) used to derive free flowing river metrics (Grill et al., 2019) and the dotted vertical line represents the limit of direct impact defined by the Environmental Impact Assessment. Lines and shaded areas are mean values and 95% confidence intervals from Generalized Additive Models (formula = y ~ s(x, bs = “cs”), family = quasibinomial) that are added as a visual aid to illustrate trends in the values. Photos illustrating dry season river level and nesting area (b) before (October 2015) and (c) after (October 2018) reservoir filling (51°36’16W, 0°53’6N, approximately 70 km upstream of the Cachoeira Caldeirão dam).

## 4 DISCUSSION

This study provided robust evidence of a reduction in *P. unifilis* nesting areas caused by the filling of a run-of-the-river dam reservoir in the Eastern Amazon. The availability of nesting areas was not related to seasonal precipitation, but was directly linked to the distance from the hydropower dam.

The present study showed that the distance to the dam has implications for the nesting patterns of *P. unifilis*. The density of nesting areas in the Control zone remained stable, whereas in the Impact zone the reduction in the density of nesting areas was pronounced after reservoir filling. Even after four years of operation, the nesting areas of *P.unifilis* in the Impact zone were not restored to levels registered for the same area in the period before dam reservoir filling (Norris, Michalski, & Gibbs, 2018a). Although it could be expected that overtime the newly formed reservoir shoreline may present suitable nesting habitat, within the most heavily impacted zone (13 - 46 km from the dam) a total of five new nesting areas have been found in the years following reservoir inundation. As the majority of new nesting areas (3/5) were found during the 2016/2017 nesting season (Norris, Michalski, & Gibbs, 2018a) the results suggest that the natural formation of new nesting areas is likely to take much longer than necessary to avoid extirpation of the impacted populations. The lack of available habitat may lead females to select poor quality nesting habitat (e.g. clay substrate) which can affect important parameters including incubation time, hatchling success and sex ratios (Girondot et al., 1998; Pignati et al., 2013). Limited habitat availability may also increase predation risk to nesting females and/or nests/eggs (Quintana et al., 2019), which may contribute to increase the extinction vulnerability of this species in the study region.

A permanent rise in dry season river levels may increase impacts of seasonal losses of nesting areas due to increasing occurrence of extreme flooding events across Amazonia (Barichivich et al., 2018) that can have negative impacts on Amazonian freshwater turtle populations that depend on terrestrial areas for nesting (Eisemberg et al., 2016; Norris, Michalski, & Gibbs, 2020). Such dry season river level increases are also likely to affect other species that use terrestrial nesting habitats. For example not only did reservoir filling flood crocodile nests at reservoirs created by two different run-of-river dams (Campos, 2019; Campos, Muniz, & Magnusson, 2019) but also appeared to affect reproduction of a female black caiman, *Melanosuchus niger* (Campos, 2019). Dam impacts do not occur in isolation from additional threats to aquatic species. In India one of the last remnant populations of another crocodilian, the Critically Endangered gharial (*Gavialis gangeticus*) may experience extirpation due to changes in availability of suitable sandbar nesting habitat (Vashistha et al., 2021). With a reduction in suitable nesting habitat across a protected but regulated river system affecting reproductive success and potentially limiting recruitment of breeding adults (Vashistha et al., 2021).

Total rainfall can impact the amount of nesting habitat available or the availability of other limiting resources, as demonstrated for *Batagur kachuga* where it had a significant negative correlation with nest number and monsoon magnitude (Sirsi et al., 2017). In contrast to Sirsi et al. (2017), the present study found that seasonal precipitation was not related to the density of nesting areas over seven years of standardised nesting area monitoring along more than 100 km of river. The lack of association with rainfall was not surprising as nesting occurs during the dry season when there is little rainfall. These seasonal differences in rainfall and flow rates have been exacerbated by deforestation across Amazonia (Stickler et al., 2013) and are one of the main reasons why Amazonian hydropower dams have failed to meet expectations in terms of power output particularly during the dry season when human energy demands reach their peak (Chaudhari et al., 2021; Fearnside, 2014, 2018). Several solutions are available, including placing turbines at lower levels, integrating with alternative energy sources such as solar power during the dry season and in-stream turbines (Chaudhari et al., 2021). Unfortunately these options are not included in current development plans. The availability of energy from renewable resources far exceeds demand in Brazil (Brasil, MME, & EPE, 2020). Although energy demand under an expansion scenario will increase from 300 to 600 million tonnes of oil equivalent by 2050 there is an annual potential of approximately 7.4 billion tonnes of energy from renewable sources (Brasil, MME, & EPE, 2020). Brazil could install 89 GW of solar power by 2050 (Brasil, MME, & EPE, 2020), but this value is likely underestimated as it did not include the majority of the 5 Mkm^2^ Legal Brazilian Amazon, which was excluded based on outdated map of areas that have not experienced human impacts. Yet, such plans do not reflect more immediate development strategies. For example a recent request from the Belo Monte dam operator (Norte Energia) to change their company bylaws to include thermo-electric power generation, which would be the first step to enable thermo-electric generation to compensate for low dry season output of the Belo Monte complex (Borges, 2019).

There is need for comprehensive and robust evaluation of local contexts to enable development of environmental flow rates that can meet dry season demands. An example from Australia showed how an increase in river levels early in the nesting season enabled water levels to remain low during incubation and hatching thereby creating a “win-win” scenario (McDougall et al., 2015). This risk assessment approach would also allow multiyear operational variation such that a consecutive sequence of multiple bad flow rate years for sandbar nesting species including turtles can be avoided. Such operational changes are challenging in Brazil as the electricity generated by hydropower dams is normally first transmitted to a centrally controlled network (“Sistema Interligado Nacional” - SIN) and then redistributed to individual States. Therefore flow rates and power outputs typically reflect national demands and are not adapted to local or regional needs. Such a separation between local impacts (supply) and increasing national demands creates inequalities for both conservation and development. For example, in November 2020 there was a 21 day power cut across Amapá state, while the three hydropower plants on the Araguari river were fully operational and transmitting electricity to the SIN. Studies are now evaluating the possibility to construct infrastructure (e.g. substations) to enable direct connection between the hydropower plants and the Amapá State electricity system (Coimbra, 2020). Such a direct link could enable simultaneous state level controls on energy generation and development of environmental flow regulation more appropriate to the state and regional conditions.

The submersion of nesting areas agrees with previous studies that suggest the impacts of run-of-river dam reservoirs are likely underestimated (Li et al., 2020; Norris, Michalski, & Gibbs, 2018a). Run-of-river dams are known to alter flow rates upstream of the reservoir, reducing flow rates which can lead to increasing upstream river levels and water residence times (Almeida et al., 2019a). Although run-of-river dams are considered to generate hydropower with relatively reduced impacts (Gaudard, Avanzi, & De Michele, 2018), due to highly seasonal rainfall and river flow rates the vast majority of Amazonian run-of-river dams include reservoirs e.g. Belo Monte (Hall & Branford, 2012) and can therefore generate drastic impacts on flowrates (Mendes et al., 2021). The results of the present study suggest that the dry season extent of river level rise is permanently at least 20 km beyond recognized limits. This finding that impacts extended beyond those of the environmental impact assessment suggests not only that caution must be taken in the calculation of reservoir extent used in greenhouse gas emissions (Almeida et al., 2019b), but also adds to the growing consensus regarding the link between superficial impact assessments and a lack of social and environmental sustainability of Amazonian hydropower developments (Fearnside, 2014, 2018; Gerlak et al., 2020).

In the case of the Cachoeira Caldeirão dam, the environmental impact assessment (EIA) of reservoir impacts was based on simplistic predictions without any consideration of uncertainty or seasonality. The spatial extent of impacts was predicted based on a single value for a “normal” flood quota of 58.3 m, not considering the range to the estimated maximum flood quota of 59.6 m (Ecotumucumaque, 2013). As such, impacts beyond those indicated in the EIA are to be expected and must be included in conservation management and mitigation actions. The comparison with global scale estimates (Grill et al., 2019) suggests that the impact of dams could be widely underestimated for biodiverse and complex tropical rivers. The analysis was based on a single species but tropical rivers include innumerable biodiversity that remains to be evaluated. Indeed, even for arguably the most intensely studied group (fishes) a recent review established a lack of results documenting impacts of run-of-river dams in tropical regions (Turgeon et al., 2021). Therefore, it is likely that the finding of impacts extending 20 km beyond the EIA limit is likely to be a minimum estimate. There is an urgent need for studies to generate more robust evidence documenting the biological impacts of hydropower developments in tropical regions.

Protected areas with the sustainable use of natural resources (e.g. IUCN Category VI) such as those around our study area allow “the sustainable use of natural resources as a means to achieve nature conservation” (IUCN, 2021). However, although protected areas are fundamental for nature conservation in the Brazilian Amazon, they often experience illegal activities, mainly related with illegal vegetation degradation, commercial fishing, hunting and mining (Kauano, Silva, & Michalski, 2017). Moreover, a previous study in our region, showed that law enforcement patrols, ordered by the protected areas management team as a nest protection strategy had little effect on illegal river turtle nest harvesting compared with community management, which was associated with a significant reduction in harvest rates of *P. unifilis* nests (Norris, Michalski, & Gibbs, 2018b). Thus, although the presence of protected areas and effective government enforcement are necessary components of conservation actions plans they are unlikely to ameliorate negative effects from hydropower developments in isolation.

## CONSERVATION RECOMMENDATIONS

A robust understanding of hydropower dam impacts is necessary to enable development of effective minimization and mitigation actions. Actions are required to not only prevent extinctions (Rhodin et al., 2018) but to also facilitate the recovery of impacted populations (Grace et al., 2021; Norris et al., 2019; Tickner et al., 2020). The nesting density of yellow-spotted river turtle (*Podocnemis unifilis*) was negatively affected by distance to a run-of-the-river dam reservoir. Hydropower developments should adopt flow regulation schemes that are compatible with the persistence of impacted species. Assessments of the spatial extents of hydropower development impacts must consider relevant range of flood quota values in order to avoid underestimation of greenhouse gas emissions and impacts to local and regional biodiversity. We suggest that mitigation actions should include not only habitat creation and restoration but also dry season flow rate regulation to increase the availability of suitable nesting areas and avoid possible negative impacts to *P. unifilis* populations at least within 70 km upstream of the Cachoeira Caldeirão dam.

Among the short- and medium-term mitigation measures that should be adopted are: i) creation of new turtle nesting habitat, with substrates suitable for the temperature dependent sex determination of this species, ii) improved regulation of dry season flow rates by the hydropower dam to enable upstream river levels to drop during the months needed for turtle nesting, incubation and hatching (September, October, November and December in the study region) and to also avoid extreme flooding events (Norris, Michalski, & Gibbs, 2020) and iii) integrated monitoring and conservation of turtle populations, particularly with the local community. It is likely that all three of these measures are required to ensure the long-term conservation and recovery of the populations in the study area (Campos-Silva et al., 2018; Norris et al., 2019; Páez et al., 2015). For example without community involvement actions such as nesting habitat creation are unlikely to be successful due to continued hunting of adults and harvest of nests in the region. Although the impacts and the recommendations listed above were shared with the Cachoeira Caldeirão operating company in 2018, years of meetings and negotiations have resulted in no interventions or financial recompense by the company to address the impacts that went beyond limits established in the Environmental Impact Assessment. The lack of conservation and restoration projects materialising as mitigation actions limits the ability of this run-of river dam to be considered as a sustainable source of energy generation.

The results show that monitoring nesting areas provides useful insight into the impacts of dams. Considering the widespread distribution of *P. unifilis* such monitoring can be widely applied to enable basin level evaluations of river level changes following dam construction. Continuous and standardized monitoring of nesting-areas can be implemented and strengthened through the establishment of community management programs (Harju, Sirén, & Salo, 2018; Rivera et al., 2021). Several studies have shown the benefits of this type of management focused on the conservation of nesting-areas and maintaining the recruitment of turtle populations (Campos-Silva et al., 2018; Norris, Michalski, & Gibbs, 2020; Norris et al., 2019). In addition, these actions bring conservation co-benefits for other species, for example migratory waterfowl, which also use the turtles’ nesting habitat (Campos-Silva et al., 2021). Similarly, local communities can also obtain economic and food benefits (Harju, Sirén, & Salo, 2018; Rivera et al., 2021).

Quantifying changes in nesting areas is important but additional studies are needed to establish a direct link with species population viability. For example, boat surveys could also provide an effective means of monitoring female populations during the nesting season as they tend to bask on logs and rocks (Conway-Gómez, Reibel, & Mihiar, 2014; Coway-Gomez, 2007). Although still incipient, studies show that boat surveys can be used to monitor freshwater turtle abundances at a local scale [e.g. 6 km surrounding islands in Florida (Breininger et al., 2019)] and more widely [e.g. *P. unifilis* populations along > 50 km of rivers in Bolivia (Coway-Gomez, 2007; Yapu-Alcázar et al., 2018) and Peru (Norris et al., 2011; Pitman et al., 2011)]. As *P. unifilis* is widespread and ubiquitous across Amazonian river basins (Norris et al., 2019; Turtle Taxonomy Working Group, 2017), integrating nesting area surveys with river based population monitoring could therefore provide robust empirical estimates to parameterize population viability analysis in basins and rivers impacted by hydropower developments across Amazonia.

Due to their social and cultural importance freshwater turtles represent conservation “flagship” species across tropical South America. Monitoring nesting areas is straight forward and should be a regional/ basin wide focus to not only rapidly identify impacts but also develop effective mitigation actions for biodiversity conservation in tropical rivers impacted by hydropower dams. Any management program to be implemented must be designed considering the social, economic and environmental characteristics of each region, as well as having the participation of the local population, research groups, government and companies in charge of dam management.

## ACKNOWLEDGEMENTS

We are grateful to Alvino Pantoja and Gilberto Souza dos Santos for their invaluable assistance during field-work in 2019. We are also grateful to the field assistants, students and volunteers who have helped to collect data during the previous nesting seasons. We would like to thank the Universidade Federal do Amapá and the Instituto Chico Mendes de Conservação da Biodiversidade (ICMBio) for providing logistical support; IBAMA for authorization (IBAMA/SISBIO permit 49632-1) to conduct research around FLONA.

## DATA AVAILABILITY

The data that supports the findings of this study are available in the supplementary information of this article. A copy of the data is also openly available via the Center for Open Science at https://doi.org/10.17605/OSF.IO/KQ573.

